# Resource-efficient pooled sequencing expands translational impact in solid tumors

**DOI:** 10.1101/2021.06.06.447265

**Authors:** Renzo G. DiNatale, Roy Mano, Vladimir Makarov, Nicole Rusk, Esther Drill, Andrew Winer, Alexander Sankin, Angela Yoo, Benjamin A. Freeman, James J. Hsieh, Ying-Bei Chen, Jonathan A. Coleman, Michael Berger, Irina Ostrovnaya, Timothy A. Chan, Paul Russo, Ed Reznik, A. Ari Hakimi

**Affiliations:** Computational Oncology Service, Epidemiology & Biostatistics Department, Memorial Sloan Kettering Cancer Center, New York, USA; Immunogenomics and Precision Oncology Platform, Memorial Sloan Kettering Cancer Center, New York, USA; Urology Service, Department of Surgery, Memorial Sloan Kettering Cancer Center, New York, USA; Biostatistics, Epidemiology & Biostatistics Department, Memorial Sloan Kettering Cancer Center, New York, USA; Department of Urology, SUNY Downstate Health Sciences University, Brooklyn, NY; Department of Urology, Montefiore Medical Center and Albert Einstein College of Medicine, Bronx, NY; Department of Medicine, University of Washington, Washington D.C., USA; Department of Pathology, Memorial Sloan Kettering Cancer Center, New York, USA; Center for Molecular Oncology, Memorial Sloan Kettering Cancer Center, New York, USA

**Keywords:** cancer genomics, cancer evolution, intratumoral heterogeneity, next-generation sequencing, somatic mutation, clonality

## Abstract

Intratumoral genetic heterogeneity (ITH) poses a significant challenge to utilizing sequencing for decision making in the management of cancer. Although sequencing of multiple tumor regions can address the pitfalls of ITH, it does so at a significant increase in cost and resource utilization. We propose a pooled multiregional sequencing strategy, whereby DNA aliquots from multiple tumor regions are mixed prior to sequencing, as a cost-effective strategy to boost translational value by addressing ITH while preserving valuable residual tissue for secondary analysis. Focusing on kidney cancer, we demonstrate that DNA pooling from as few as two regions significantly increases mutation detection while reducing clonality misattribution. This leads to an increased fraction of patients identified with therapeutically actionable mutations, improved patient risk stratification, and improved inference of evolutionary trajectories with an accuracy comparable to *bona fide* multiregional sequencing. The same approach applied to non-small-cell lung cancer data substantially improves tumor mutational burden (TMB) detection. Our findings demonstrate that pooled DNA sequencing strategies are a cost-effective alternative to address intrinsic genetic heterogeneity in clinical settings.

## MAIN TEXT

Clear cell renal cell carcinoma (ccRCC), the most common and aggressive form of kidney cancer, is characterized by extensive intratumoral heterogeneity (ITH) whereby driver mutations frequently arise only in a subset of tumor cells [1–3]. As a result of ITH, clinically informative but subclonal mutations are commonly missed by the standard practice (at our institution [4] and others [5]) of sequencing single tumor regions. In a landmark multiregional sequencing study of 101 ccRCC tumors, the TRACERx consortium reported that fifty-six percent of all detected mutations were subclonal [6], and ∼20% of subclonal mutations had demonstrable clinical value either for prognostication in clinical risk models (*TP53, BAP1*, and *PBRM1*) [7], or as criteria for administration of targeted therapy (*MTOR, TSC1*, and *PTEN*) [8] (**Figure 1a)**. Single region sequencing places a hard constraint on the sensitivity to detect and study mutations for two reasons: somatic mutation dropout (*i*.*e*. absence of a mutation in the particular tumor region sampled) and erroneous clonality assertions (*i*.*e*. attributing mutations as clonal when in fact they are only subclonal, or *vice versa*). Multiregional sequencing strategies address ITH by sequencing the genomic material of several spatially-separated regions of the same tumor[9]. However, due to the added sequencing expenses, this approach becomes prohibitively costly as the number of regions increases, limiting its use in practice.

**Figure 1.**
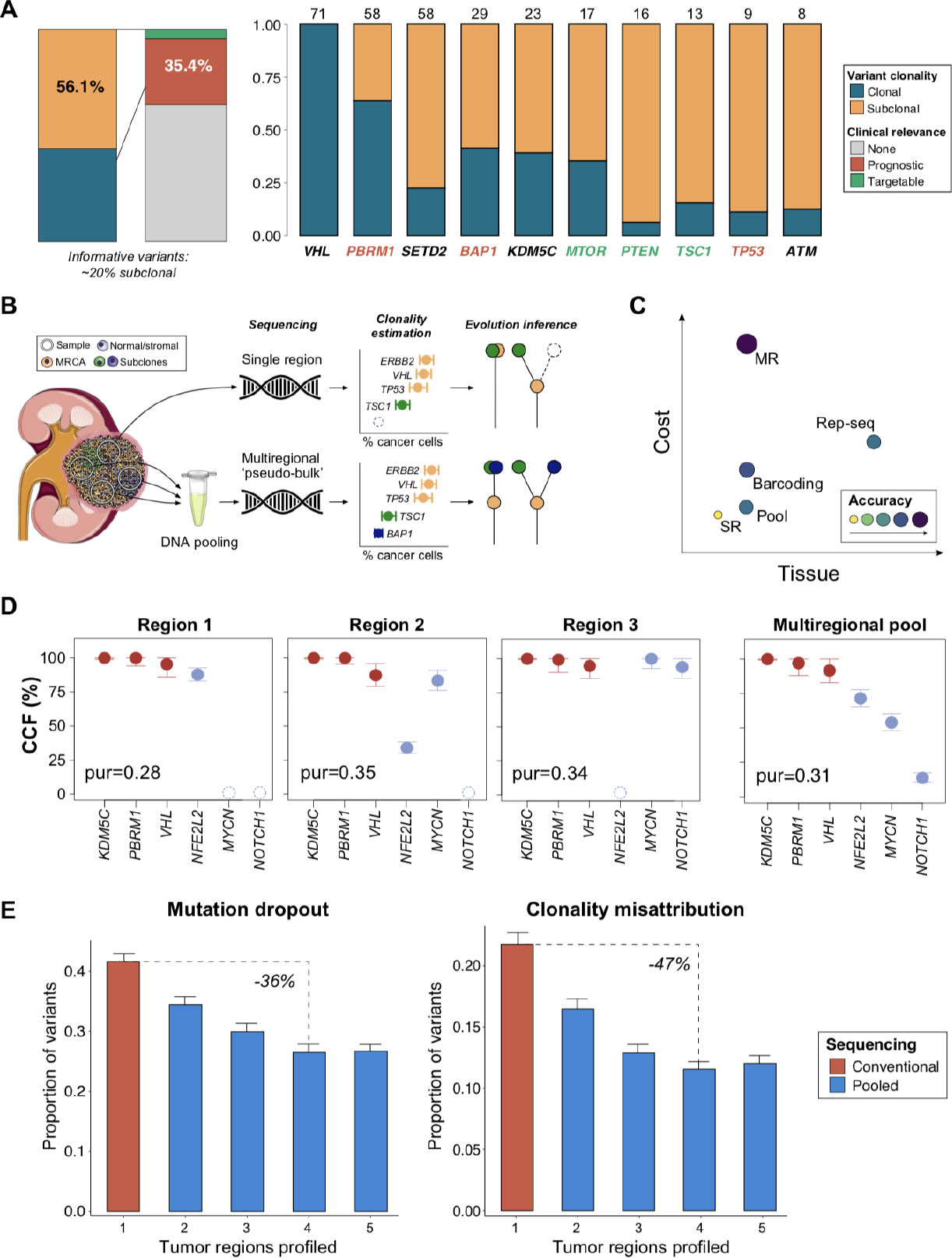
Intratumoral mutational heterogeneity in renal cell carcinoma can be overcome with pooled sequencing. **A**.Variants identified in the TRACERx RCC cohort. The clonal status and proportion of clinically-relevant variants are shown (left). Top 10 most commonly-mutated genes in the TRACERx RCC cohort, by clonal status and clinical relevance (right). The numbers at the top represent the number of unique variants identified per gene in the cohort. **B.** Schematic representing the confection of tumor DNA pools. During ‘Evolution inference’, evolutionary trees are depicted with (right) and without (left) spatial resolution. **C**. Schematic comparing resource requirements and accuracy across different sequencing approaches. MR: conventional multi-regional sequencing, SR: single-region sequencing. **D**. Variant cancer-cell fraction (CCF) identified in three separate tumor regions and its corresponding multiregional DNA pool. Results are shown in percentages relative to the total number of cancer cells in the sample. **E**. Dropout (left) and clonality misattribution estimates (right) from the *in silico analysis* performed on the TRACERx RCC cohort are shown for conventional and pooled sequencing (red and blue bars, respectively). Average event-level results are shown with their 95% confidence intervals (error bars) calculated across 100 simulation

We reasoned that a more cost-effective approach to quantitatively managing ITH would be to pool samples from many regions together into a single “pseudo-bulk” before library construction (**Figure 1b)**. Doing so would potentially ameliorate mutation dropout by increasing the likelihood of capturing a subclonal mutation, while reducing the mis-attribution of clonal status to mutations present only in single regions of the tumor. The benefits of a pooled approach would come at several costs: first, from diluting the sequencing bandwidth devoted to each individual region, and second, from loss of spatial information that would be obtained from *bona fide* multiregional sequencing. However, complete loss of spatial information could potentially be avoided (with an increase in cost and overhead) by barcoding DNA libraries before sequencing, an approach that has previously been demonstrated by several investigators[10]. Furthermore, pooling of tumor regions preserves precious tumor tissue which could be used for further molecular, immunohistochemical, or other profiling, and therefore is a material-efficient alternative to fully unbiased representative sequencing [11]. Direct pooling of DNA samples thus represents a flexible, cost-efficient middle ground strategy that can be readily implemented into current pipelines without requiring additional expertise or reagents (**Figure 1c)**.

We examined the feasibility of pooled sequencing using a deep, targeted clinical sequencing platform. For each of six ccRCC tumors, six spatially discrete regions were selected and pooled into a single sample. In parallel, we sequenced separate aliquots of the same tumor regions to standard depth, generating a ground-truth set of variant calls (**Figure 1d** and **Supplementary figure 1a**). One case (RCC006) had no variants identified in any region and was not included in the mutational analysis (**Supplementary table 1, 2)**. Multiregional pooled sequencing of six regions at an average depth of ∼900x (150x/region) resulted in a mutation dropout rate of 4.3% (1/23 variants) and a clonality error rate of 4.5% (1/22 variants) (**Supplementary figure 2a**). Compared to single region profiling, pooled multi-regional sequencing showed a 12% lower dropout rate (95% CI: 2.0 - 22.4%, Welch *t*-test, p=0.02) and a 13% lower clonality error rate (95% CI: 1.2 - 24.9%, Welch *t*-test, p=0.03) (**Supplementary Figure 2b)**. Reduction in clonality misattribution was robust with the chosen cancer-cell fraction (CCF) threshold, with an estimated Matthew’s correlation coefficient (MCC) of 0.73 (with +1 indicating perfect prediction and −1 complete disagreement) **(Supplementary figure 2c, d)**. Notably, all the mutations missed/misclassified were present in the tumor with the highest regional variability in purity (RCC004, with a variance 5-fold higher than the average, *σ*^*2*^= 0.05 vs 0.01), and none of the pooled samples had purity estimates below our quality threshold (compared to 14%, or 5/36, of the regions profiled separately). These findings demonstrate an additional potential advantage of pooled sequencing, *i*.*e*. the possibility to reduce sample failure rates during clinical sequencing.

To validate our findings and further assess the utility of this sequencing strategy, we analyzed multiregional sequencing data from the TRACERx consortium. The validation cohorts consisted of 101 individuals with a ccRCC diagnosis (median, 8 tumor regions [range, 2-75]) profiled with a sequencing panel targeting 110 cancer genes at a median depth of 612x (range, 105–1,520x) (TRACERx RCC cohort, [6]), and 100 patients with non-small cell lung cancer (TRACERx NSCLC cohort), profiled with exome sequencing at a median depth of 431x (range 83-986x) for tumor regions and 415x (range 107-765x) for the matched germline (median, 3 tumor regions, range 2-8) [12] **(Supplementary table 3)**. From our own data, we confirmed that tumor purity estimates in DNA pools were predictable *in silico* to high accuracy using tumor purity from single regions (**Supplementary figure 2e)** Next, we simulated pooled sequencing in the TRACERx data (at equivalent depth to single-region sequencing) using a bootstrapping procedure (**see Methods**). Outcomes were then calculated on each random sample and averaged to produce region-number-specific estimates.

Pooled sequencing substantially decreased mutation dropout relative to single region profiling, even with the addition of just a single region (17% decrease in dropout with a pool of two regions). Similarly, we observed that this approach significantly improved our ability to correctly assign clonality to observed mutations, with a 24% drop in clonality assignment error with the addition of a single region to a pool (**Figure 1e)**. When evaluating these same outcomes at the patient-level (i.e. proportion of individuals with at least one variant dropped/misclassified), we observed that pooled sequencing with a single additional region would result in a 14% decrease in both the number of patients subject to mutation dropout and the number affected by misattribution of clonality **(Supplementary figure 3b, d)**. Consistent with the rarity of spatially-delimited low-allele-frequency mutations in ccRCC (arising in cancer genes), we observed a negligible number of false-negative mutation calls with higher number of regions **(Figure 2a)**. No differences were observed between tumor pools of four regions and those with higher numbers when evaluating their reliability when attributing mutation clonality, however, this result was found specific to the tumor type context **(Supplementary figure 4a, b)**. Importantly, because the financial cost of next-generation sequencing (NGS) assays is dominated by sequencing costs (*i*.*e*. related to library size due to depth and breadth) rather than sample processing and genomic material extraction, obtaining DNA from multiple regions and mixing them into a single pseudo-bulk would result in minimal additions to the total cost **(Supplementary Figure 5a)**. Given that our direct pooling approach requires no additional reagents (nor modifications to the computational infrastructure), it occupies a flexible middle ground compared to *bona fide* multiregional sequencing and full-scale mixing of left-over tumor tissue [10,11]. We defined a metric of cost-effectiveness as the change in mutation dropout (or clonality) per tumor relative to the change in cost (**Supplementary figure 5 b, c**). Using cost estimates for targeted panel sequencing from our own institution, we compared the cost-effectiveness of conventional and pooled multi-regional sequencing relative to single-region profiling. Pooled sequencing (of 2 to 4 tumor regions) was found to be ∼10% more cost-effective than single-region profiling both for mutation detection and clonality assessment, while the opposite was observed with conventional multi-regional sequencing. Notably, the added benefit of pooled sequencing was lost when pooling 10 regions or more **(Figure 2b)**. We next examined the translational utility of pooling discrete regions of a tumor in the management of ccRCC by several metrics. In patients with metastatic ccRCC receiving first-line treatment with tyrosine kinase inhibitors, the mutation status (irrespective of clonal status) of *PBRM1, BAP1*, and *TP53* is of prognostic significance [7], and dropout of somatic variants in these genes therefore affects risk stratification. We observed that risk stratification would be affected in 10% of the TRACERx RCC patients if only a single region were sequenced. Pooled sequencing of 4 regions corrected the risk stratification in 4% of patients, effectively reducing the baseline error in risk stratification by 22% **(Figure 2c)**. Furthermore, the presence of mutations in a subset of genes represent potential therapeutically actionable targets and/or eligibility criteria for clinical trials. Pooled sequencing (of 4 regions) significantly increased the number of patients identified with such mutations by more than 70% (from 6% to 10%) (**Figure 2d)**.

**Figure 2.**
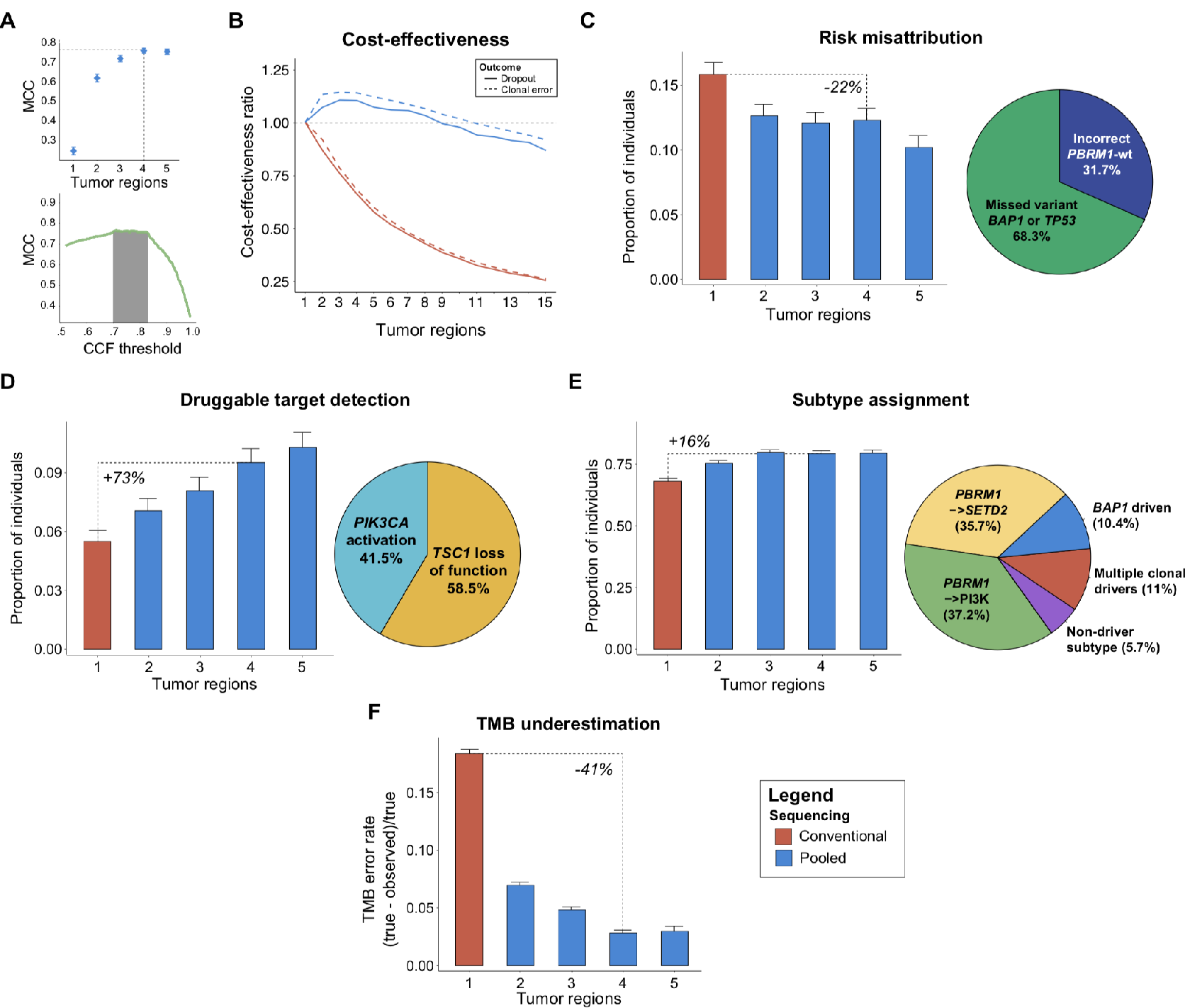
Translational value of pooled sequencing. **A.** MCC estimates used to evaluate the optimal number of regions to pool (top) and the optimal range of CCF thresholds to define clonality (bottom). **B**. Cost-effectiveness analysis of the dropout and clonality error rates (at the tumor-level) between conventional and pooled multiregional assessment. Results are shown for increasing numbers of tumor regions relative to the cost-effectiveness of a single tumor region (ratio=1). **C**. Proportion of patients subject to risk misattribution (left). The most-common features resulting in risk misattribution after pooling 4 regions were dropout of *BAP1*or *TP53* variants, followed by dropout of *PBRM1* variants (right). **D**. Proportion of patients with at least one targetable mutation identified (left). The most-commonly missed targetable alterations in pools of 4 regions were *TSC1* loss-of-function and *PIK3CA* activating mutations (right). **E**. Proportion of patients in which the correct molecular subtype was determined based on mutational data (left). The most-commonly missed subtypes in a 4-region tumor pool are shown in the pie chart (right). **F**. Underestimation of tumor mutational load. Bars represent the average TMB error across all simulations (i.e. true - sample / true) and error bars its 95%CI. *TMB:* tumor mutational burden, *MCC:* Matthew’s correlation coefficient, *CCF:* cancer-cell fraction.

Independent of its translational value, pooled sequencing provides a cost-effective lens onto patterns of ITH. Recent work by the TRACERx consortium in ccRCC has proposed “evolutionary subtypes” based on the presence and clonality of mutations in five genes (*VHL, PBRM1, SETD2, BAP1 and PTEN*). We examined our capacity to correctly assign evolutionary subtypes in pooled sequencing according to the heuristics outlined by Turajlic and colleagues [6]. Pooling four tumor regions increased the correct evolutionary subtype assignment by 16%, with the majority of missed subtypes corresponding to the ‘*PBRM1*→PI3K’ and ‘*PBRM1*→*SETD2’*subtypes, with relatively good outcomes (**Figure 2e**). Pooled sequencing thus represents a potential strategy for the interrogation of subclonal mutational diversity and inference of evolutionary trajectories, which have further implications for patient outcomes.

Finally, we explored the translational value of pooled sequencing in the context of an entirely different disease and sequencing platform. In a cohort of 100 NSCLC patients from the TRACERx consortium, we evaluated *in silico* the utility of pooled sequencing in accurately quantifying tumor mutation burden (TMB); this measure is employed as a biomarker for response to immunotherapy in this disease. While single region sequencing underestimates total tumor mutation burden by nearly 20%, the addition of a single region to a DNA pool reduced this effect by 41% (**Figure 2f)**. Since the clonality of neoantigens is an emerging determinant of T cell immunoreactivity [13], and given that TMB and neoantigen load are highly correlated [14], accurate assessment of mutation burden with multiregional approaches may improve prognostication in the context of immunotherapy for NSCLC. However, the ability of a clonality-aware TMB measure to predict response to immune-checkpoint blockade will need to be evaluated in this context, as it is currently optimized to the single region setting[15].

Intratumoral heterogeneity is a fundamental hurdle in the genomically-informed delivery of care to cancer patients. In ccRCC, such heterogeneity is so pervasive that it confounds the accurate identification of the small set of driver mutations of therapeutic relevance. Our proposed approach of direct pooled DNA sequencing from several tumor regions overcomes some of these issues at a fraction of the cost of *bona fide* multiregional profiling; and it does so without excess use of precious tissue material, preserving it for subsequent profiling studies. Pooling thus represents a viable and cost-effective strategy to overcome ITH during clinical sequencing. Importantly, the overhead costs for both single region and pooled sequencing, including sample acquisition, data handling, storage, and analysis, are largely the same, as the size of the sequencing library remains identical. Furthermore, multiregional DNA pooling allows for the inclusion of additional processing steps before sequencing, providing an extra degree of flexibility when adjusting this approach to different clinical scenarios. Finally, by mixing regions of variable purity, we also envision that pooled sequencing may ameliorate the ∼3% of tumor samples (∼300/10,000 per year total) which currently fail clinical sequencing at our institution due to excessively low tumor purity, thus increasing resource utilization efficiency [16].

Although our current analysis is limited to single nucleotide variants and indels, copy number variants (CNVs) could be similarly evaluable by pooled sequencing. However, the relatively low density of heterozygous SNPs tiling the genome in targeted sequencing platforms renders the attribution of clonality to CNVs extremely challenging[9]. One might speculate that ongoing refinement of targeted sequencing panels or the use of broader panels could create new opportunities for copy-number analysis from pooled sequencing.

## DATA AVAILABILITY

All the processed data and statistical code needed to reproduce the findings in this study have been provided in the supplementary materials as well as in a publicly-available repository (https://github.com/reznik-lab/DNApooling_RD). The raw sequencing data (MSK-IMPACT) produced in this study are deposited on the Sequence Read Archive (SRA) under the accession number PRJNA633220. Data from the validation sets are available in the supplementary materials of the original TRACERx publications [6,12], only the filtered/annotated versions are provided with this manuscript. Any other relevant data is available from the corresponding authors upon reasonable request.

## ACKNOWLEDGMENTS

We thank Charles Swanton, Samra Turajlic, Nicholas McGranahan, Kevin Litchfield and all the members of the TRACERx consortium for facilitating the mutation data used in the study. We thank the Reznik and Chan Lab members for helpful discussions. We thank the staff and physicians of the MSK Department of Medicine Kidney Program and the Urology Service for helpful suggestions. We acknowledge the use of the Integrated Genomics Operation Core, funded by the NIH/NCI Cancer Center Support Grant (CCSG, P30 CA008748), Cycle for Survival, and the Marie-Josée and Henry R. Kravis Center for Molecular Oncology. This work was in part supported by grants NIH R35 CA232097 and DOD KC180165 (TAC), as well as the DOD Kidney Cancer Research Program W81XWH-18-1-0318 and the Kidney Cancer Association Young Investigator Award. (ER).This work was also supported by the Mellnikoff Fund (TAC) and the Weiss Family Fund (AAH).

## AUTHOR CONTRIBUTIONS

*Sample and patient data procurement:* AW, AS, YC, PR, JC, AY, BAF

*Data processing:* VM, RGD, MB, TAC

*Statistical analysis:* RGD, ED, IO, ER

*Manuscript preparation:* RGD, RM, NR, ER, AAH

*Oversight*: AAH, ER, TAC, PR, JH

## SUPPLEMENTARY MATERIALS

**Supplementary table 1.** Characteristics of the samples profiled

| Patient | Gender | Age | Tumor regions | Race | T stage | Clinical stage | Overall stage | Tumor size (cm) | Nuclear grade | Histologic subtype |
| --- | --- | --- | --- | --- | --- | --- | --- | --- | --- | --- |
| RCC001 | Male | 69 | 6 | Caucasian | pT3a | cT3a | III | 6 | 3 | Clear cell renal cell carcinoma |
| RCC002 | Female | 42 | 6 | Asian | pT1b | cT1b | I | 4.2 | 2 | Clear cell renal cell carcinoma |
| RCC003 | Male | 53 | 6 | Caucasian | pT3a | cT1b | III | 4 | 4 | Clear cell renal cell carcinoma |
| RCC004 | Male | 37 | 6 | Caucasian | pT3a | cT3aN0M1 | IV | 9.3 | 4 | Clear cell renal cell carcinoma |
| RCC005 | Male | 74 | 6 | Caucasian | pT3a | cT1bN0M1 | IV | 7.5 | 3 | Clear cell renal cell carcinoma |
| RCC006* | Male | 61 | 6 | Caucasian | pT3b | cT3aN1M0 | III | 9.5 | 4 | Clear cell renal cell carcinoma |

\* No nonsynonymous somatic mutations were identified in any of the samples profiled from this tumor

– **Supplementary table 2**. Mutation-annotation file of samples profiled with MSK-IMPACT sequencing as part of this study

**Supplementary table 3.** Clinicopathologic characteristics of the individuals in the TRACERx cohorts.

|  | levels | RCC (n=101) | NSCLC (n=100) |
| --- | --- | --- | --- |
| **Tumor regions profiled (number)** | median [IQR] | 8 [5, 15] | 3 [2, 4] |
| **Age (years)** | median [IQR] | 64 [57, 69] | 68 [63, 76] |
| **Sex (%)** | Female | 33 (32.7) | 38 (38.0) |
|  | Male | 68 (67.3) | 62 (62.0) |
| **Ethnicity (%)** | Caucasian | 57 (56.4) | 97 (97.0) |
|  | Afrocaribbean | 37 (36.6) | 3 (3.0) |
|  | Asian | 7 (6.9) | 0 (0.0) |
|  | Unknown | 0 (0.0) | 0 (0.0) |
| **Smoking status (%)<sup>a</sup>** | Never smoked | 44 (43.6) | 12 (12.0) |
|  | Ex-smoker | 28 (27.7) | 48 (48.0) |
|  | Smoker | 12 (11.9) | 40 (40.0) |
|  | Unknown | 17 (16.8) | 0 (0.0) |
| **Histology (%)<sup>b</sup>** | Clear cell renal cell carcinoma | 101 (100.0) | 0 (0.0) |
|  | Adenocarcinoma | 0 (0.0) | 61 (61.0) |
|  | Squamous cell carcinoma | 0 (0.0) | 32 (32.0) |
|  | Other | 0 (0.0) | 7 (7.0) |
| **Stage (%)<sup>c</sup>** | T1 | 30 (29.7) | 62 (62.0) |
|  | T2 | 7 (6.9) | 24 (24.0) |
|  | T3 | 56 (55.4) | 14 (14.0) |
|  | T4 | 8 (7.9) | 0 (0.0) |
| **Vascular invasion (%)<sup>d</sup>** |  | 54 (53.5) | 41 (41.0) |
| **Adjacent organ invasion (%)<sup>e</sup>** |  | 8 (7.9) | 27 (27.0) |
| **Distant metastasis (%)<sup>f</sup>** |  | 24 (23.8) | 0 (0.0) |
| **Multifocal renal tumor (excluded) (%)<sup>f</sup>** |  | 5 (5.0) | 0 (0.0) |

a. Individuals listed as 'recent ex-smokers' in the NSCLC cohort were considered smokers
b. Category 'Other' includes the following NSCLC histologies: large cell carcinoma, large cell neuroendocrine carcinoma, carcinosarcoma, adenosquamous carcinoma
c. The NSCLC cohort only included resectable patients (i.e. stages 1 through 3)
d. For RCC tumors includes any of the following types of invasion: microvascular, renal vein or inferior vena cava
e. For RCC tumors this represents invasion Beyond Gerota's fascia (i.e. stage T4), while in NSCLC patients it represents pleural invasion
f. Only applies to RCC tumors

– **Supplementary table 4**. Mutation-annotation file of the TRACERx RCC cohort annotated with cell abundance estimates.
– **Supplementary table 5**. Mutation-annotation file of the TRACERx NSCLC cohort annotated with cell abundance estimates.

**Supplementary figure 1.**
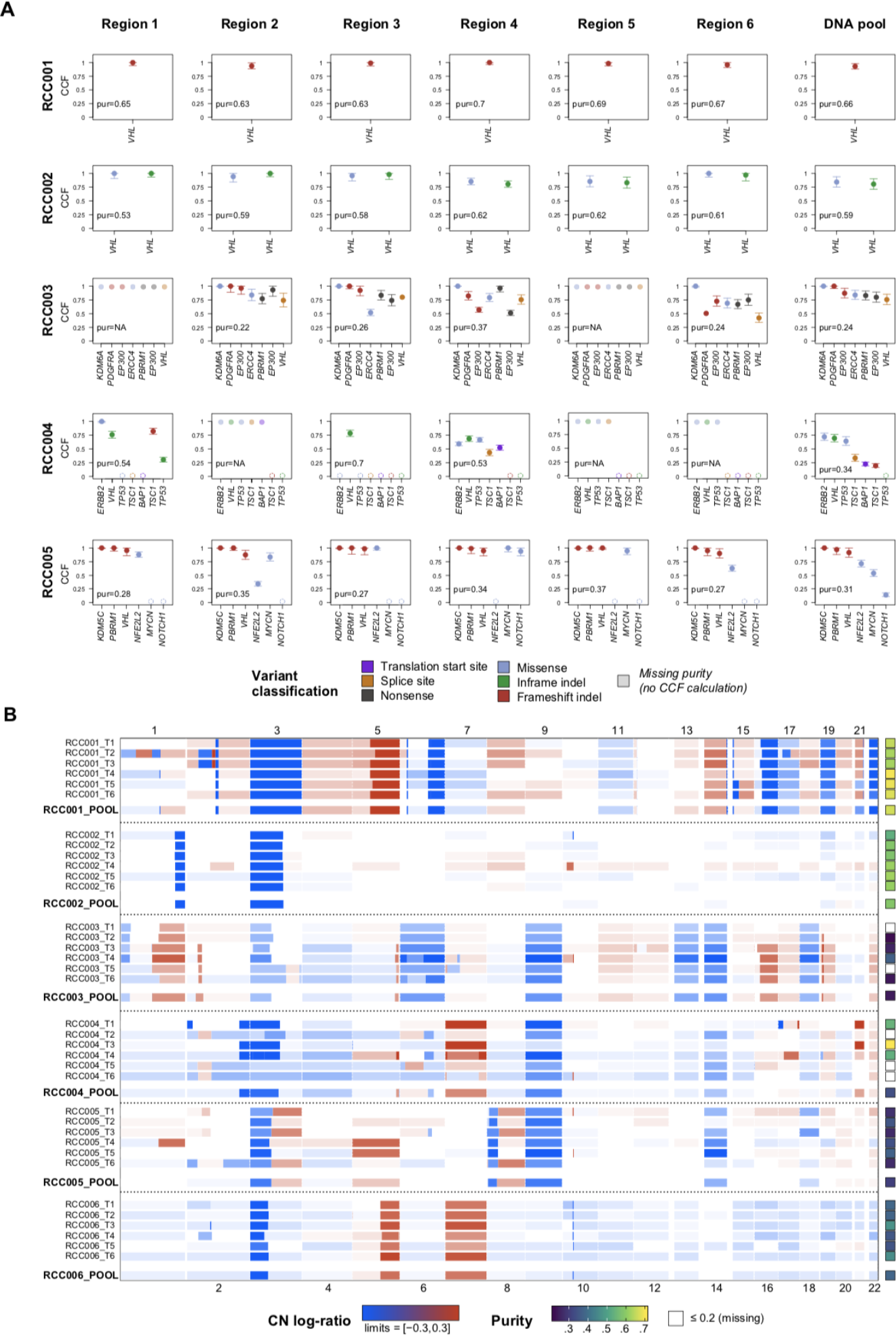
Detailed mutation and copy-number calls seen in the samples profiled. **A.** Mutation cancer-cell fraction (CCF) estimates in each of the tumor samples sequenced. Six spatially-separated regions and a DNA pool were profiled for each tumor, and no mutations were detected in any of the samples from RCC006 (not shown). In samples where purity could not be estimated, the variants detected are shown at CCF=1 (dimmed out). Mutations that were not detected are shown at CCF=0 (dashed circles). **B**.Multi-regional and pooled copy-number (CN) profiles observed in the different samples. The heatmap shows the gains (red) and losses (blue) evidenced across the genome of each tumor region. The tiles on the right represent the purity calculated in each sample. Purity values below 20% are not estimated by the algorithm and appear as missing (white).

**Supplementary figure 2.**
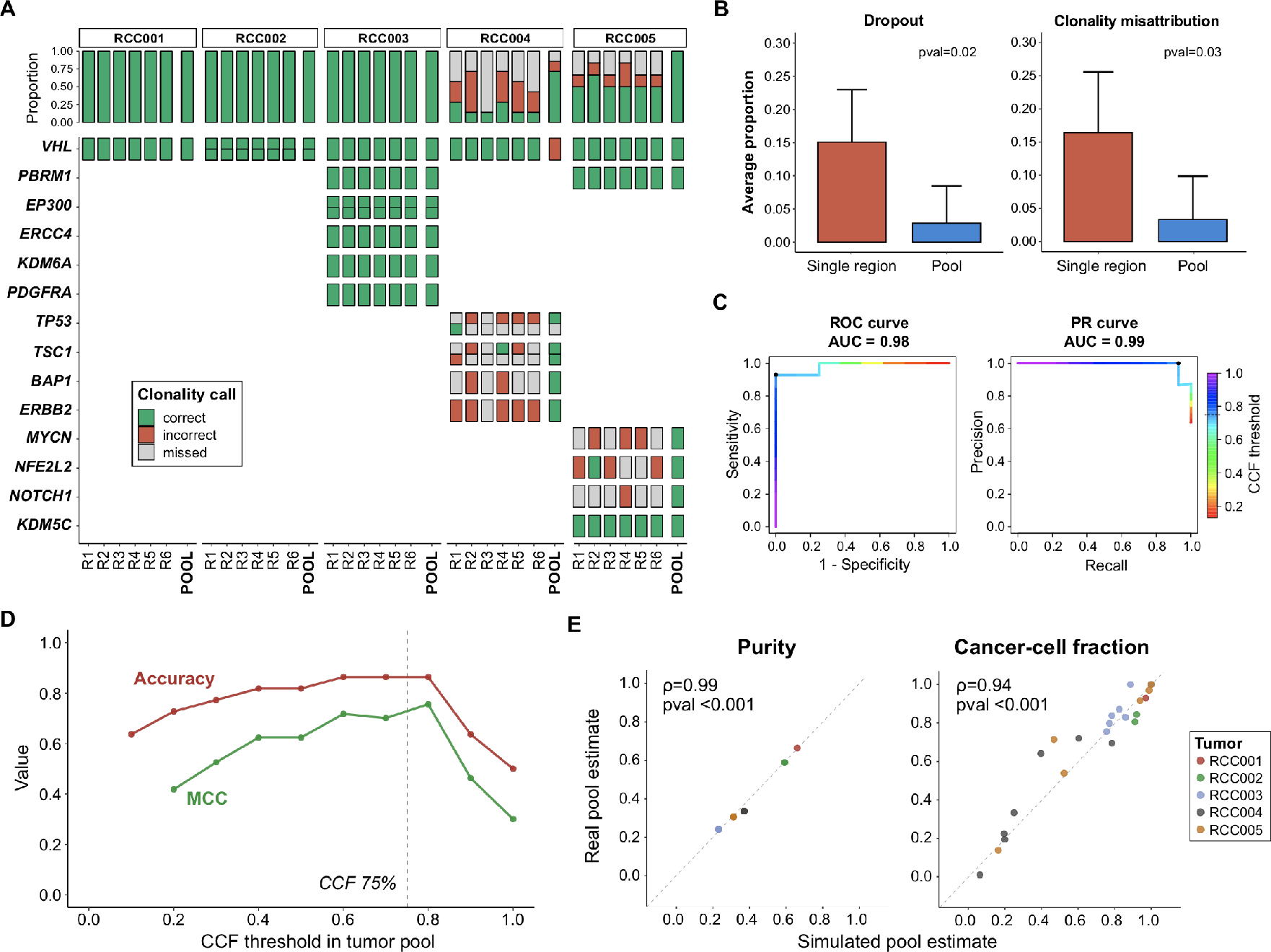
Comparison of mutation dropout and clonality misattribution between pooled and single-region sequencing using targeted sequencing. **A.**Oncoprint showing the nonsynonymous variants identified in the samples (N=23). The color of the tiles represent the concordance (green) or discordance (red) between the true clonality status (from assessment of all the regions individually) and the assertion made based on evaluating each sample individually (MR threshold: 0.5, POOL threshold: 0.75, **see Methods, Clonality assessment**). Samples not detected are shown in grey. **B**. Average proportion of variants dropped (left) and misclassified (right) in the samples profiled, comparing tumor DNA pools and single regions. Significant differences (Welch’s two-sample *t*-test) were observed with both outcome measures. **C**. Receiver-operating characteristic (ROC) and precision-recall (PR) curves to classify mutation clonality using the CCF values in confected DNA pools. The line colors represent the CCF threshold. The color of the lines represents the CCF range and the black dots represent the threshold used to define clonality in the study. **D**. Performance of tumor DNA pools to attribute mutation clonality across the range of all possible CCF thresholds. Colored lines represent different classifier performance measures (accuracy and MCC). The dashed line represents the threshold of 75% CCF used in the study. **E**. Observed versus predicted cell-abundance estimates. The purity (left) and cancer-cell fraction (CCF, right) values from the confected DNA pools are compared to those estimated by assessing multiple separate regions (Spearman-rank correlation test). *CCF:* cancer-cell fraction, *MCC:* Matthew’s correlation coefficient.

**Supplementary figure 3.**
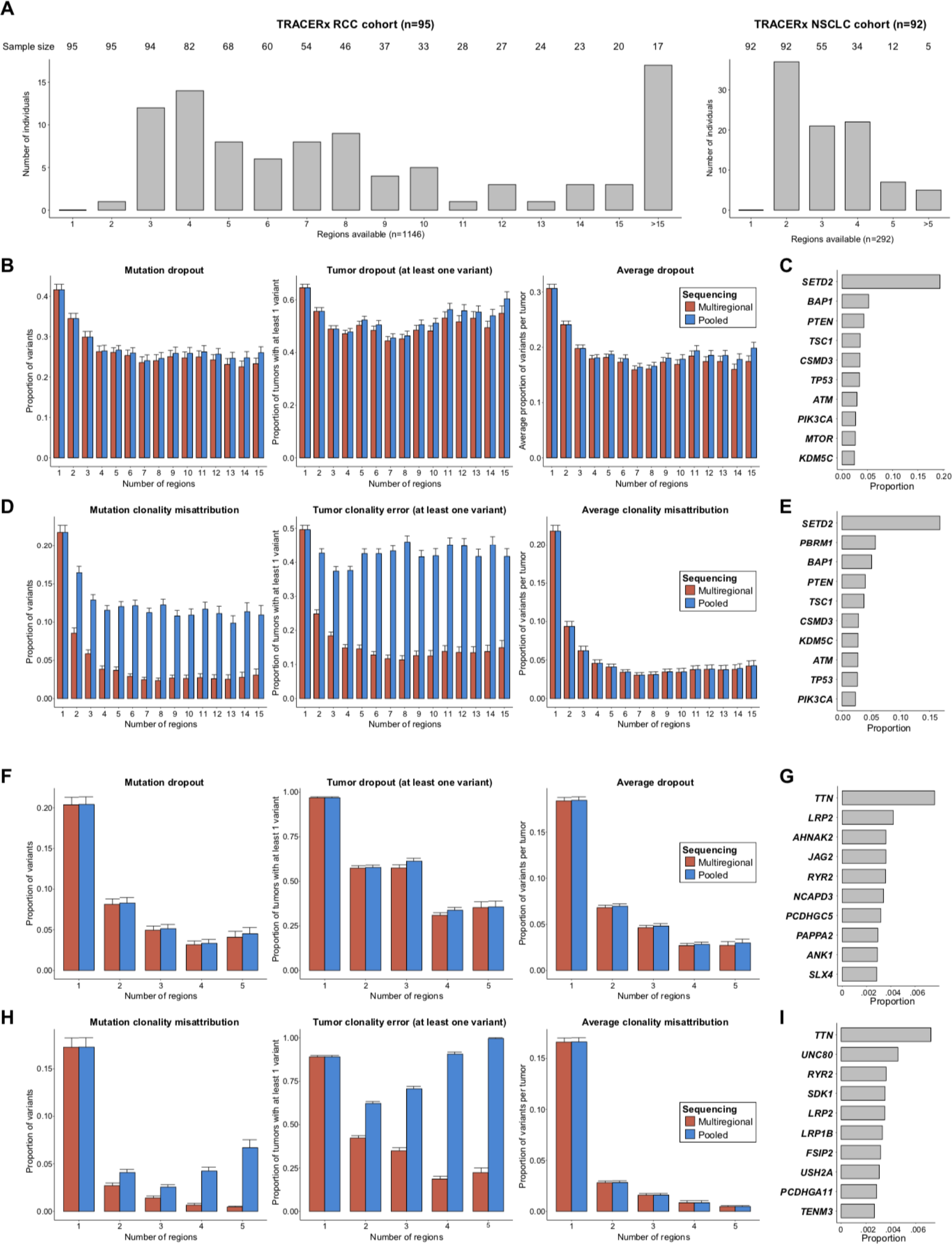
Dropout and clonality misattribution outcomes in the TRACERx cohorts. **A**.Number of individuals included in the analysis by regions available. Results are shown separately for the RCC (left) and NSCLC (right) cohorts. Numbers at the top represent the total number of tumors with at least x regions that were used to calculate sample sizes in the bootstrapping approach. **B.** Mutation dropout in the RCC cohort. Three separate outcomes are listed (for B,D,F,H): total proportion of variants dropped (left), total proportion of patients with at least one variant dropped (middle), and average proportion of variants dropped per tumor (right). Red and blue bars represent the average estimates (over 100 bootstrapping iterations) and the error bars their 95% CI. **C**. Top 10 genes dropped across 100 simulated pools of 4 regions. Results are expressed relative to all the variants dropped. **D**. Clonality misattribution in the RCC cohort. **E**. Top 10 genes misclassified across 100 simulated pools of 4 regions. Results are expressed relative to all the variants misclassified. **F**. Mutation dropout in the NSCLC cohort. **G**. Top 10 genes dropped across 100 simulated pools of 4 regions. Results are expressed relative to all the variants dropped. **H**. Clonality misattribution in the NSCLC cohort. Top 10 genes misclassified across all simulated pools of 4 regions. Results are expressed relative to all the variants misclassified.

**Supplementary figure 4.**
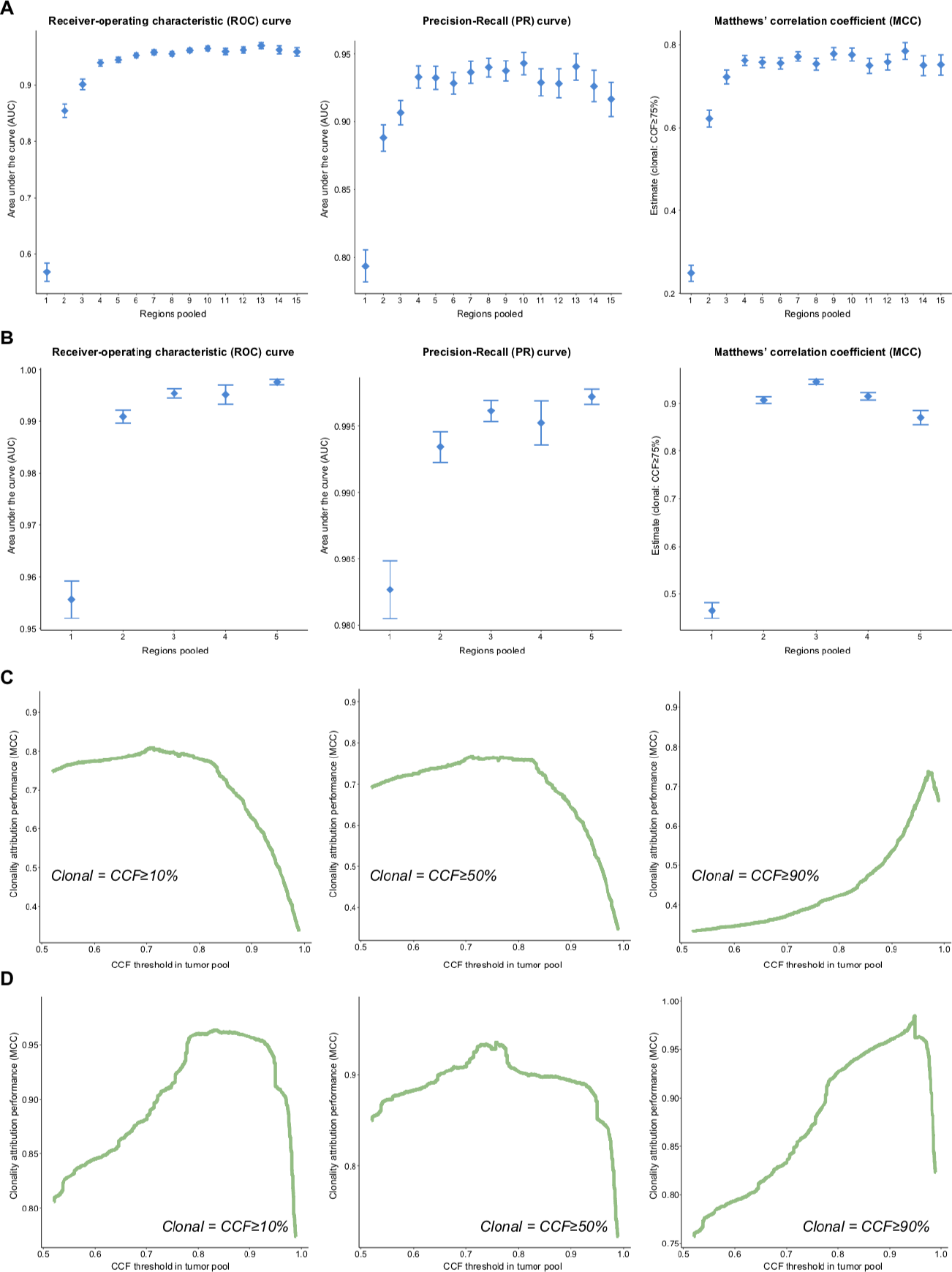
Performance of pooled sequencing CCF estimates to attribute mutation clonality. **A.**Binary classifier performance by number of regions in the RCC cohort. Area under the curve estimates for the ROC curve (left) and PR curve (middle), as well as the MCC (right), are shown. The MCC was calculated using a CCF threshold of 0.75. Average estimates (across 100 bootstrapping iterations) along with their 95% confidence intervals are shown. **B**. Binary classifier performance by number of regions pooled in the NSCLC cohort (as above, A). This cohort had a lower number of regions overall. **C**. Pooled CCF estimate performance with different definitions of ‘true clonality’. Average MCC estimates from all simulated pools of 4 regions are shown. Each panel represents a different CCF threshold used to define true clonality with all the regions available(shown in italics). Clonal variants were defined as those present in every region at a CCF equal or higher to that threshold. **D**. CCF estimates performance in the NSCLC cohort (as above, C). *ROC:* Receiver-Operating Characteristic, *MCC:* Matthews’ Correlation Coefficient, *PR:* Precision-Recall curve, *RCC:* renal cell carcinoma, *NSCLC:* non-small-cell lung cancer, *CCF:* cancer-cell fraction.

**Supplementary figure 5.**
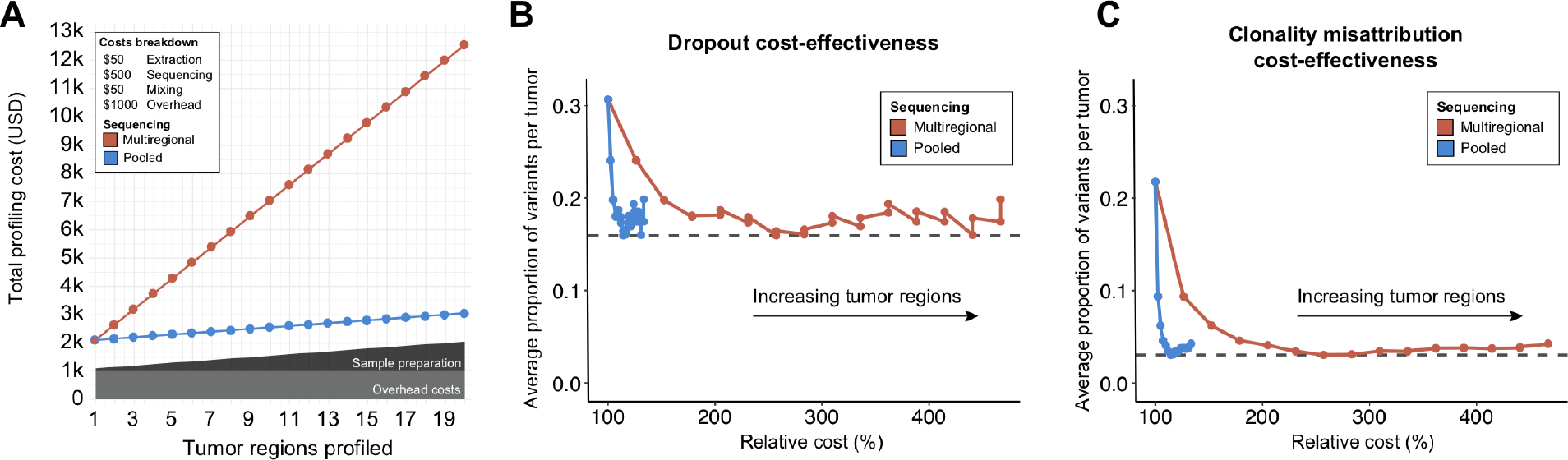
Cost-effectiveness of pooled and multi-regional sequencing with increasing tumor regions. **A.**Total costs of profiling with different sequencing strategies. Results are broken down by expense and shown in absolute numbers (1000 USD = 1k). **A**. Average dropout rate per tumor and cost increments relative to single-regions. **B**. Average clonality misattribution rate per tumor and r cost increments relative to single-regions.

– **Supplementary material 1**. Estimation of relative cellular abundances in pooled tumor samples using multi-region information
– **Supplementary material 2**. Script containing the code in R for the comparison of single-versus multi-region pooled sequencing in the *in-situ* experiment.
– **Supplementary material 3**. Script containing the code in R for the bootstrapping procedure performed in the TRACERx cohorts.
– **Materials and methods**

### METHODS

#### Patients and sample procurement

Tumor tissue and matched-blood from six kidney cancer patients who underwent partial or radical nephrectomy at our institution were obtained as follows. Peripheral venous blood samples were drawn in the clinic during pre-surgical visits and used as matched germline controls. Primary tumor specimens were procured during the surgical procedure and six *ex-vivo* core-needle biopsy samples were obtained from spatially-separated regions. All samples were snap-frozen and stored until genomic material extraction. Specimens were also reviewed by a genitourinary pathologist (Y.C.) and a diagnosis of clear cell renal cell carcinoma was established in all cases. Median tumor size (maximum diameter, in cm, on pathologic review) was 6.75 cm (range, 4 to 9.5) and five of the tumors were pT3 stage (**Supplementary table 1**). One of the tumors (RCC006) did not show any non-silent mutations in any of the regions and was excluded from further analysis. The final sample size consisted of 30 tumor regions from five patients as well as five tumor DNA pools (each one containing DNA from six regions).

#### Ethics

The institutional ethics review board pre-approved all research-related activities. All individuals provided written informed consent for the participation of this study, which included their tumor and normal tissue being profiled and their clinical data being shared in a non-identifiable manner. All activities related to this study were conducted in accordance with the principles of Good Clinical Practice and the Declaration of Helsinki.

#### DNA extraction and multiregional pool construction

Samples from each tumor region were stored at −80*C for 3-4 weeks. After thawing the samples in regular ice, DNA extraction was performed using the DNeasy Blood & Tissue Kit according to the manufacturer’s protocol (QIAGEN catalog #69504). Genomic DNA from tumor regions and blood samples was eluted in 50 ml of nuclease-free water. Tumor DNA pools were confected by combining 10 ml of the eluate from each region and mixing them by pipetting into a single 60 ml sample. A total of 8 DNA samples from each patient (1 blood, 6 tumor regions and 1 tumor pool) were sent for sequencing at the Integrated Genomics Operations Core of Memorial Sloan Kettering Cancer Center, a CLIA-certified laboratory (see Acknowledgments).

#### Sequencing, raw data processing and somatic mutation calling

Sequencing was done using MSK-IMPACT^®^, a targeted sequencing panel approved by the FDA for the study of solid tumors. It captures the exons of 341 cancer genes as well as a set of known single-nucleotide polymorphisms (SNPs) tiled across the genome (for CN estimation)[4]. The median sequencing depth for tumor and normal samples was 876x (range, 574x - 1911x) and 544x (range, 500x - 802x), respectively. Raw sequencing data in FASTQ format was aligned to the human genome (version b37) using the Burrows–Wheeler aligner (BWA v.0.7.10). Local realignment was performed with the Genome Analysis Toolkit (GATK v4.1.4.1[17]) and duplicate reads removed using Picard v.2.13.

For somatic variant calling, we used our integrated bioinformatics pipeline which includes four different variant calling tools: MuTect2 (part of GATKv4.1.4.1[17]), Strelka2 v2.9.10[18], Varscan v2.4.3 [19] and Platypus [20]. Ancillary filters were used to obtain high-accuracy calls, these included: a coverage of at least 10x in the tumor, with 5 or more reads supporting the variant, a variant allele frequency (vAF, i.e. proportion of DNA read) greater or equal than 5% in single-region and 2% in pooled tumor samples, and a vAF below 2% in the matched normal sample. Only nonsynonymous somatic mutations were considered, and single-nucleotide variants (SNVs) identified at a frequency higher than 1% in dbSNP[21] or 1000Genomes project[22] were excluded. Finally, dubious calls were manually reviewed by investigators using the Integrative Genome Viewer software v2.4.10 for additional accuracy [23,24]. A total of 23 unique nonsynonymous somatic mutation events were considered in the analysis. Tumor mutational burden (TMB) was defined as the total number of mutations identified in a tumor specimen (considering all available regions).

#### Allele-specific copy-number (ASCN) analysis

Allele-specific copy number analysis (ASCN) was performed using FACETS v0.5.6 [25] through a publicly-available R package (https://github.com/mskcc/facets/). By comparing the tumor and copy-number (CN) profiles, this tool provides integer copy number (CN) values for each tumor allele in addition to other genome-wide estimates, such as tumor ploidy and sample purity (i.e. the proportion of DNA in tumor samples derived from tumor cells).

#### Clonality assessment: cancer-cell fraction (CCF) estimation

We used the cancer-cell fraction (CCF) of each somatic variant as a surrogate of the relative timing of each mutation. This measure represents the proportion of tumor cells bearing a somatic variant of interest, and has been extensively used as a clonality surrogate in several landmark cancer genomic studies[26–28]. The CCF of each variant was calculated assuming a diploid CN state (i.e. total CN=2, minor allele CN=1) in the normal tissue (CPN_norm_) and using the vAF, locus-specific ploidy (CPN_mut_) and sample purity (p) information as described by McGranahan et al. [29]:

#### Cell abundance estimates in pooled DNA samples

We derived the formulas to estimate sample purity and mutation CCF values in a hypothetical tumor DNA pool using multi-regional data (**Supplementary Material 1b**). Although the expected purity of a tumor pool is a near-average of the value in each region, CCF cannot be estimated the same way. Owing to the fact that total DNA from different tumor samples was combined, and that CCF represents a measure of mutation abundance relative to the cancer cell compartment (rather than all cells), single-region CCF averages are not appropriate to estimate the values in the pool. Therefore, we considered the mutant cell fraction (MCF, i.e. proportion of mutant cells, out of all cells) and purity of each region as follows:

#### TRACERx analyses

We obtained publicly-available somatic mutation data from the original RCC and NSCLC studies of the TRACERx consortium. Individuals who met the following criteria were included in the analysis: i) absence of bilateral or multifocal renal tumors, and ii) more than one primary tumor region with non-silent mutations available. The final RCC cohort included 92 individuals (1146 regions, profiled with a sequencing panel targeting 110 cancer genes), while the NSCLC cohort included 95 individuals (292 regions) profiled using exome sequencing **(Supplementary figure 3a)**. Mutation-annotation files (MAF) containing the CCF and the cell-abundance estimates used in the analysis were provided (**Supplementary table 4, 5**).

#### Somatic variant annotation

Nonsynonymous exonic somatic mutation data both from the in-house samples and TRACERx cohorts were parsed into MAF format using the maf2maf function (v2.4) from a publicly-available package (https://github.com/mskcc/vcf2maf). The MAFs were then annotated with the corresponding CCF data using an R implementation of the previously-described formula (https://github.com/mskcc/facets-suite). Finally, the evidence on the treatment implications of each mutation was assessed using OncoKB [8] through its Python implementation (https://github.com/mskcc/facets-suite). Only levels of evidence 1 through 3 were included in the analysis, and the database was queried on March 6^th^ of 2020.

#### Clonality definitions

Clonality definitions in sequencing studies vary widely and different methods and definitions have been used in single and multi-regional sequencing [9]. Most commonly, specific thresholds are set in the CCF continuum to classify mutations into ‘clonal’ or ‘subclonal’, with some investigators opting for a third ‘indeterminate’ category. We opted for a binary classification approach using the same clonality definitions from the original TRACERx Renal publications (i.e. mutations had to be present in every region at a CCF greater or equal to 50% to be considered clonal) [6,30]. Similarly, a threshold of CCF≥75% was used to define clonality in the tumor DNA pools **(Supplementary Figure 2d and 4b)**.

Only variants at or above 5% CCF were reported in the TRACERx studies and therefore included in the analyses. To simulate subclonal mutation dropout due to excessive dilution of low frequency variants in DNA pools, we set a detection limit threshold in the simulated pools. Variants predicted to have a CCF below 2% in the tumor DNA pools were considered ‘missed’ in the analysis and counted as dropped (not considered in clonality analysis).

#### Bootstrapping procedure

To compare the outcomes of conventional and pooled sequencing, a bootstrapping procedure with 100 iterations was performed using nonsynonymous somatic mutation data from the TRACERx consortium. Analyses were conducted separately in the RCC and NSCLC cohorts due to differences in the breadth of sequencing. Briefly, given a number of tumor regions *R*, we obtained a random sample (with replacement) containing 70% of the tumors with at least *R* number of regions profiled (**Supplementary Figure 3a)**. We then created a ground-truth set of variant calls from all the regions available for these specimens and, mimicking random tissue sampling, selected a random subset of *R* regions to analyze. The variants detected and the clonality assertions made using only the chosen regions were compared to the previously-established ground truth (*i*.*e*. using all available regions); this was done both conventionally (i.e. multiple separate regions assessed) and simulating a tumor DNA pool (i.e. containing DNA from *R* regions mixed in equal proportions). Next, a series of variant- and tumor-level outcomes are calculated in each random subset of tumors, and estimates from the 100 iterations are averaged. This approach was repeated with an increasing number of regions and the estimates compared between conventional and pooled sequencing using parametric statistical testing.

#### Outcomes and measures

##### Mutational dropout and misattribution of clonality

We used a combination of variant- and tumor-level measures to compare the conventional and pooled sequencing approaches. Results are shown with an increasing number of regions in both real and simulated DNA pools **(Figure 1, 2)** as well as a comparable multiregional assessment of the same number of regions **(Supplementary Figure 3)**. In all the analyses, a ground truth set of variant calls was created using all the tumor regions available. True clonal mutations were defined as those with an estimated CCF≥0.5 in every region of the tumor. After listing all the mutations in a tumor and their true clonal status, we compared the calls to the results obtained from each sequenced sample (*in situ*) or a set of regions selected at random from the original data (*in silico*). Only mutations detected by the sample(s) of interest were considered when evaluating the clonality assertions made with both approaches.

The proportion of variants dropped and misclassified in each sample sequenced in-house was calculated, the sample-specific estimates were then averaged and compared between the individual regions and pools. For the *in silico* analysis, after obtaining the ground-truth variant calls from all the tumors in a given iteration of the bootstrapping, we calculated three measures for each of the outcomes: i) the absolute proportion of variants dropped/misclassified in all the pools selected (event-level), ii) the proportion of tumors with at least one variant dropped/misclassified (tumor-level) and, iii) the average proportion of variants dropped/misclassified per tumor (tumor-level average) (Supplementary Figure 3).

##### Performance evaluation of pooled sequencing

Next, we evaluated the performance of the CCF values estimated from pooled samples as a binary classifier of mutation clonality. Each mutation was assigned a true ‘clonal’ or ‘subclonal’ status, and these assertions were contrasted to the CCF estimates obtained from the pooled samples using the aforementioned definitions. We calculated three different estimates (at the event-level) to assess the reliability of CCF in a pooled sequencing context; results are shown as average estimates (and their 95% CIs) across the 100 iterations of the bootstrapping. To evaluate the performance of a pooled CCF value without setting a specific threshold, area under the curve (AUC) estimates were calculated for the Receiver-Operating Characteristic (ROC) and Precision-Recall (PR) curves, using the R package ‘PRROC’. Given the frequent presence of unbalanced confusion matrices in this context (only branched or only truncal mutations present), we selected the Matthew’s Correlation Coefficient (MCC) to evaluate the performance of pooled CCF estimates as binary clonality classifiers [31]. The MCC estimates are shown across all possible CCF thresholds, and results were contrasted to the definitions used in the study (i.e. CCF≥0.5 in each region for ground-truth and CCF≥0.75 in pooled samples).

##### Cost-effectiveness

The cost-effectiveness of pooled sequencing and *bona fide* multiregional sequencing were compared as follows **(Figure 2b)**. First, the total costs of profiling were calculated with both approaches using estimates from our own institution **(Supplementary Figure 5a)**. Then, the average dropout and clonality misattribution estimates were calculated across the 100 simulated samples **(Supplementary Figure 5b, c)**. Cost-effectiveness was defined as the average proportion of variants detected/correctly classified per tumor divided by the total cost of profiling. Finally, results were expressed relative to single-region profiling (relative cost-effectiveness).

##### Risk misattribution rate

To assess the translational value of the proposed approach, we evaluated the ability of pooled sequencing to correctly attribute risk points to each tumor, using the genomic criteria described by Voss et al. [7]. A total of three genes were evaluated to risk-stratify patients:

1. Presence of a *BAP1* or *TP53* mutation - 1 point
2. Absence of *PBRM1* variants, or co-occurrence with one of the above - 1 point

Each tumor was assigned a ‘true risk score’ (of maximum two points) using all the regions available, and these were contrasted to the assertions made with the regions of interest. The rate of risk misattribution was defined as the proportion of tumors in the sample with erroneous risk scores (regardless of direction).

##### Molecular subtype assignment (according to TRACERx Renal)

We evaluated the ability of pooled sequencing to correctly assign a molecular subtype to each tumor as described by Turajlic et al. [6]. Since one of the subtypes is characterized by presence of copy-number alterations, which were not evaluated as part of this study (PBRM1-->sCNA), minor adjustments had to be made in the way tumors were classified. Therefore, we could only achieve a concordance of 96% with the original results. First, a list of core driver genes (*VHL, PBRM1, SETD2, BAP1 and PTEN)* and PI3k pathway genes (*PIK3CA, MTOR, PTEN, TSC1, TSC2*) were obtained from the original study. Next, a set of rules were applied in hierarchical order to each tumor based on the mutations detected and their clonality estimates. If none of the rules were met, tumors were assigned to the ‘Non-driver’ (or indeterminate) sub-group. The rules were applied as follows:

i. ***Multiple clonal driver:*** presence of ≥ 2 *BAP1, PBRM1, SETD2* or *PTEN* clonal mutational events. *If not, then:*
ii. ***BAP1-driven:*** presence of a *BAP1* mutational driver event, and no other “core” mutational drivers in the same clone/subclone (other than *VHL*). In pooled samples, tumors were assigned to this category if they had a clonal *BAP1* variant or a subclonal one, with the highest CCF among all the drivers present (if any other were present). *If not, then:*
iii. ***PBRM1***→***SETD2:*** presence of a *PBRM1* mutation followed by a *SETD2* one. For tumor pools this meant a higher estimated CCF for the *PBRM1* variant compared to the one on *SETD2. If not, then:*
iv. ***PBRM1***→***PI3k:*** presence of a *PBRM1* mutation followed by a PI3k pathway variant. For tumor pools this meant a higher estimated CCF for the *PBRM1* variant compared to the one on the PI3k pathway gene(s). *If not, then:*
v. ***PBRM1***→***CNA*:** presence of a *PBRM1* mutation followed by a driver somatic CNA. Since the CNAs were not evaluated, tumors were put tentatively in this category when a *PBRM1* variant was observed without any other mutational drivers (except for *VHL*). However, if the tumor met the criteria for the ‘*VHL* wild-type’ subtype, they were assigned that group instead. *If not, then:*
vi. ***VHL wild-type:*** absence of *VHL* mutation. *If not, then:*
vii. ***VHL monodriver:*** *VHL* as the only “core” driver mutation.

##### TMB error (underestimation) rate

TMB estimates were only reported for the NSCLC TRACERx cohort due to the low number of genes assessed in the RCC cohort (∼110), which precludes accurate TMB assessment, as well as the lack of clinical value in this setting [32]. Differences in TMB detection were compared between conventional and pooled sequencing by calculating an average TMB error estimate across the 100 simulated cohorts. For each patient, this was calculated by subtracting the TMB from a single region or a pool (of >1 region) to the ‘true TMB’ (i.e. the sum of all unique events), and the result expressed relative to the ground truth (i.e. true - observed/true).

#### Statistical analyses

Outcomes were compared between different sequencing approaches using parametric statistical testing. Average estimates were compared between conventional and pooled sequencing using *t*-tests. Unpaired tests were used given the different samples or random sets of patients used in the *in-situ* and *in-silico* analyses, respectively. Pearson correlation tests were used to compare observed versus expected cell-abundance estimates in the pooled samples sequenced in this study. Sensitivity analyses were performed to assess the performance of pooled CCF estimates for attributing clonality to mutations across different conditions. First, we explored the performance of CCF estimates of tumor pools to classify clonality using increasing tumor regions **(Supplementary figure 4a, b)**. Next, we tested the effect of changing the definitions of true clonality. This was done by setting increasing CCF thresholds to define ‘true clonality’ in the conventional multi-regional assessment used as reference (i.e. CCF≥10%, 50% and 90%, respectively, **Supplementary Figure 4c, d**). Results were reported as point estimates along with their 95% confidence intervals (i.e. averages across 100 simulations). Hypothesis tests were always two-sided and statistical significance defined as a p-value below 0.05. All analyses were performed using the R platform v3.5.3.

